# Prior Uncertainty Impedes Discrete Locomotor Adaptation

**DOI:** 10.1101/2023.08.27.554999

**Authors:** Aojun Jiang, Francis M. Grover, Mary Bucklin, Jasjit Deol, Anna Shafer, Keith E. Gordon

**Author notes:** Corresponding Author: Francis M. Grover, 645 N Michigan Ave, Suite 1100, Chicago, IL 60611. A. Jiang and F. M. Grover contributed equally to this work.

## Abstract

The impact of environmental uncertainty on locomotor adaptation remains unclear. Environmental uncertainty could either aid locomotor adaptation by prompting protective control strategies that stabilize movements to assist learning or impede adaptation by reducing error sensitivity and fostering hesitance to pursue corrective movements. To explore this, we investigated participants’ adaptation to a consistent viscous force field after experiencing environmental uncertainty in the form of unpredictable balance perturbations. We compared the performance of this group (Perturbation) to the adaptive performance of a Control group that did not experience any unpredictable perturbations. Perturbations were delivered using a cable-driven robotic device applying lateral forces to the pelvis. We assessed whole-body center of mass (COM) trajectory (COM signed deviation), anticipatory postural adjustments (COM lateral offset), and first step width. The Perturbation group exhibited larger disruptions in COM trajectory (greater COM signed deviations) than the Control group when first walking in the force field. While the COM signed deviations of both groups decreased towards baseline values, only the Control group returned to baseline levels. The Perturbation groups COM signed deviations remained higher, indicating they failed to fully adapt to the force field before the end. The Perturbation group also did not adapt their COM lateral offset to counter the predictable effects of the force field as the Control group did, and their first step width increased more slowly. Our findings suggest that exposure to unpredictable perturbations impeded future sensorimotor adaptations to consistent perturbations.

## Introduction

Humans use a combination of neural control mechanisms to produce stable movements in new or changing environments. Different control mechanisms are typically adopted in response to either consistent or unpredictable environments (Morasso et al., 2014; Osu, Burdet, Franklin, Milner, & Kawato, 2003; Saha & Morasso, 2012). When learning to move in a consistent environment (e.g., a velocity dependent force field), humans form sensorimotor memories of the external forces that are used to plan future movements that will specifically compensate for the expected forces (Bucklin, Wu, Brown, & Gordon, 2019; Bucklin, Brown, & Gordon, 2023; Franklin, Osu, Burdet, Kawato, & Milner, 2003; Shadmehr & Mussa-Ivaldi, 1994). In contrast, when moving in a novel, unpredictable environment (e.g., random perturbations), humans adopt general protective movement strategies to compensate for external forces with high uncertainty (Bucklin, Deol, Brown, Perreault, & Gordon, 2023; Burdet, Osu, Franklin, Milner, & Kawato, 2001; Gribble, Mullin, Cothros, & Mattar, 2003; Wei, Wert, & Körding, 2010). While sensorimotor caution facilitates stability while moving in highly unpredictable environments, it is otherwise inefficient as it can produce slow, inflexible, and fatiguing movements (Hogan, 1985) and is therefore not ideal for predictable situations where predictive control is feasible.

It is unclear the degree to which exposure to environmental uncertainty impacts future adaptation to predictable environments. Uncertainty could inhibit adaptation of predictive strategies by inducing lasting decreases in error sensitivity, i.e., a cautious strategy of deliberately overlooking trial-to-trial errors (Albert et al., 2021; Fernandes, Stevenson, & Körding, 2012; Havermann & Lappe, 2010; Herzfeld et al., 2014; Therrien, Wolpert, & Bastian, 2018). This strategy could potentially persist even in consistent, predictable environments where trial-to-trial errors are useful for sensorimotor adaptation. Thus, exposure to an unpredictable environment could suppress future adaptation to a consistent environment—sensorimotor adaptation slows, or movement errors are sustained.

On the other hand, environmental uncertainty might provide a benefit to the early adaptation stage of predictive strategies by prompting protective strategies that stabilize movements while the neural commands for feed-forward movements are developed (Franklin et al., 2003). Then, as learning progresses and predictive movements become tuned to the environment, reliance on protective strategies will be reduced (Darainy & Ostry, 2008; Heald, Franklin, & Wolpert, 2018; Koji, 2007).

The notion of exposure to environmental uncertainty having a beneficial (or detrimental) impact on locomotor adaptation would have important implications for clinical rehabilitation settings. Connecting findings in upper limb reaching models (e.g., Burdet et al., 2001; Heald et al., 2018; Osu et al., 2003) to walking scenarios could yield valuable applications to gait rehabilitation, and many studies suggest similarities in the control of walking and reaching. Like reaching, investigations into trial-by-trial locomotor adaptation to a novel, consistent environment reveal large initial movement errors (Emken & Reinkensmeyer, 2005) and after-effects following practice (Lam, Anderschitz, & Dietz, 2006; Noble & Prentice, 2006), indicating the sensorimotor adaptation of predictive control (see also Blanchette & Bouyer, 2009; Choi & Bastian, 2007; Emken, Benitez, & Reinkensmeyer, 2007). However, walking differs from reaching in that it has greater consequences for failure (falls) and greater risk of failure (regular periods of instability in the gait cycle), which may result in greater priority placed on sensorimotor caution than adaptation (Bruijn & van Dieën, 2018). Thus, it is not clear if prior exposure to unpredictable environments will favorably accelerate sensorimotor adaptation or inhibit it when walking in a novel and consistent environment.

### 1.1 Current study

The current study builds upon a prior study (Bucklin, Wu, Brown, & Gordon, 2019) that observed strategies were adopted by participants to stabilize balance as they practiced stepping to a target through the novel force field. When the force field was unexpectedly removed (i.e., during catch trials), participants exhibited large whole-body movement errors in the opposite direction of the force field, indicating that their adopted strategies were largely *predictive*, i.e., they involved feedforward control in anticipation of the force field (as opposed to merely reacting to it). The force field in that study was consistent; its properties never changed from trial to trial. For the current study, we considered if the adaptations of participants were dependent on the expectation of this consistency and whether participants conditioned beforehand by uncertainty (in the form of fundamentally unpredictable forces) might experience enhanced or impeded adoption of predictive strategies to the consistent force field.

To explore this question, participants stepped quickly from a static standing position to a target located two steps ahead while experiencing disruptive forces delivered to their pelvis by a cable-driven robotic system. For the main training portion of the experiment, these forces were the same as those in the previous study (Bucklin et al., 2019): a laterally-directed viscous force field with consistent (and therefore learnable) properties. However, immediately prior to a force field block of trials, participants experienced forces with unpredictable (and therefore unlearnable) properties, including direction (left or right), timing, and number of applications during each stepping trial. We assessed walking performance and associated control strategies during the force field block relative to baseline (i.e., unobstructed walking) as well as during catch trials, which were sporadically occurring force field trials where the force field was not delivered. We then compared these assessments to those of the participants in the previous study (Bucklin et al., 2019) who experienced the force field training period without any prior experience with unpredictable forces (i.e., a control group).

### 1.2 Hypotheses

Our hypotheses were derived from observations in our previous study that, with repetitive practice, people developed predictive control strategies to preserve their center-of-mass (COM) trajectories against the same novel and consistent force field used in the current study (Bucklin et al., 2019). One of these strategies was to very quickly (i.e., within the first few trials) widen the lateral placement of their first step to help counteract the forces imposed by the force field. A second strategy was to increase the lateral shift of their COM (i.e., an anticipatory postural adjustment used to position their COM over their stance limb before taking their first step), which served to bias the participant’s initial posture in the opposite direction of the force field so that the force field would pull them back towards their desired trajectory instead of pulling them away from it. In the current study, we explored how these strategies (and overall performance) were affected for this walking task by prior exposure to environmental uncertainty created via unpredictable forces.

We considered two competing hypotheses based on the previously discussed possibilities of how inducing sensorimotor caution via prior exposure to environmental uncertainty could subsequently impact the development of predictive control and overall adaptation to the force field. The first hypothesis (**Hypothesis 1)** was that the development of predictive control would be inhibited compared to the control group, expressed as 1) smaller (or undetectable) changes in lateral first step width and COM lateral offset, 2) greater deviations in COM trajectory remaining after training, and 3) smaller errors during catch trials. The rationale for this prediction is that prior exposure to environmental uncertainty would produce lasting reductions in error sensitivity (cf. Herzfeld et al., 2014), prompting the brain to ignore motor errors and hesitate to pursue trial-to-trial movement corrections and resulting in less overall adaptation to the force field.

The competing hypothesis **(Hypothesis 2)** was that prior exposure to environmental uncertainty would improve development of predictive control compared to the control group, expressed as 1) greater changes in lateral first step width and COM lateral offset, 2) smaller deviations in COM trajectory remaining after training, and 3) greater deviations in COM trajectory during catch trials. The rationale for this prediction was that environmental uncertainty could prompt protective strategies that stabilize movements in the early trials of the main training period (cf. Heald et al., 2018). The early movement stability provided by the protective strategies would facilitate the development of predictive strategies, at which point the protective strategies would be abandoned. We predicted that this would result in greater overall adaptation to the force field.

## Methods

### 2.1 Participants

Twenty-six healthy young adults (14 females, 12 males, 23.2 ± 3.3 years of age, 67.8 ± 8.6 kilograms of body weight, 1.71± 0.07 meters of height, mean ± SD) participated in the experiment. Demographic characteristics were similar between experimental groups. All participants provided informed written consent, and all protocols were approved by the Northwestern University Institutional Review Board. Data from thirteen participants were collected as part of a previous study (Bucklin et al., 2019). All participants were free of any musculoskeletal, neural, and/or vestibular pathologies affecting gait or balance and were able to walk continuously for 30 minutes without undue fatigue or health risk. No participants had any prior experience walking in the external force fields used in the experiment.

### 2.2 Experimental setup

Participants performed a series of discrete goal-directed stepping trials. Each participant began each trial in a static standing position with their feet positioned inside a 0.3 x 0.3 m square start target projected on the walking surface (Hitachi America, Ltd). Then, following an auditory cue, the participant stepped quickly from the start target to an end target (a second 0.3 x 0.3 m square) (Fig. 1a). The distance between targets was adjusted for each participant to be 1.5x their leg length (0.896 ± 0.053 m; measured from the greater trochanter to the floor), which resulted in roughly a two-step task for all participants. For safety, participants wore a trunk harness attached to an overhead support device (Aretech LLC, Ashburn, VA) that moved freely on a passive trolley in the fore-aft direction. The shoulder straps of the trunk harness were adjusted so that they did not provide bodyweight support or restrict lateral movements during the trials.

**Figure 1.**
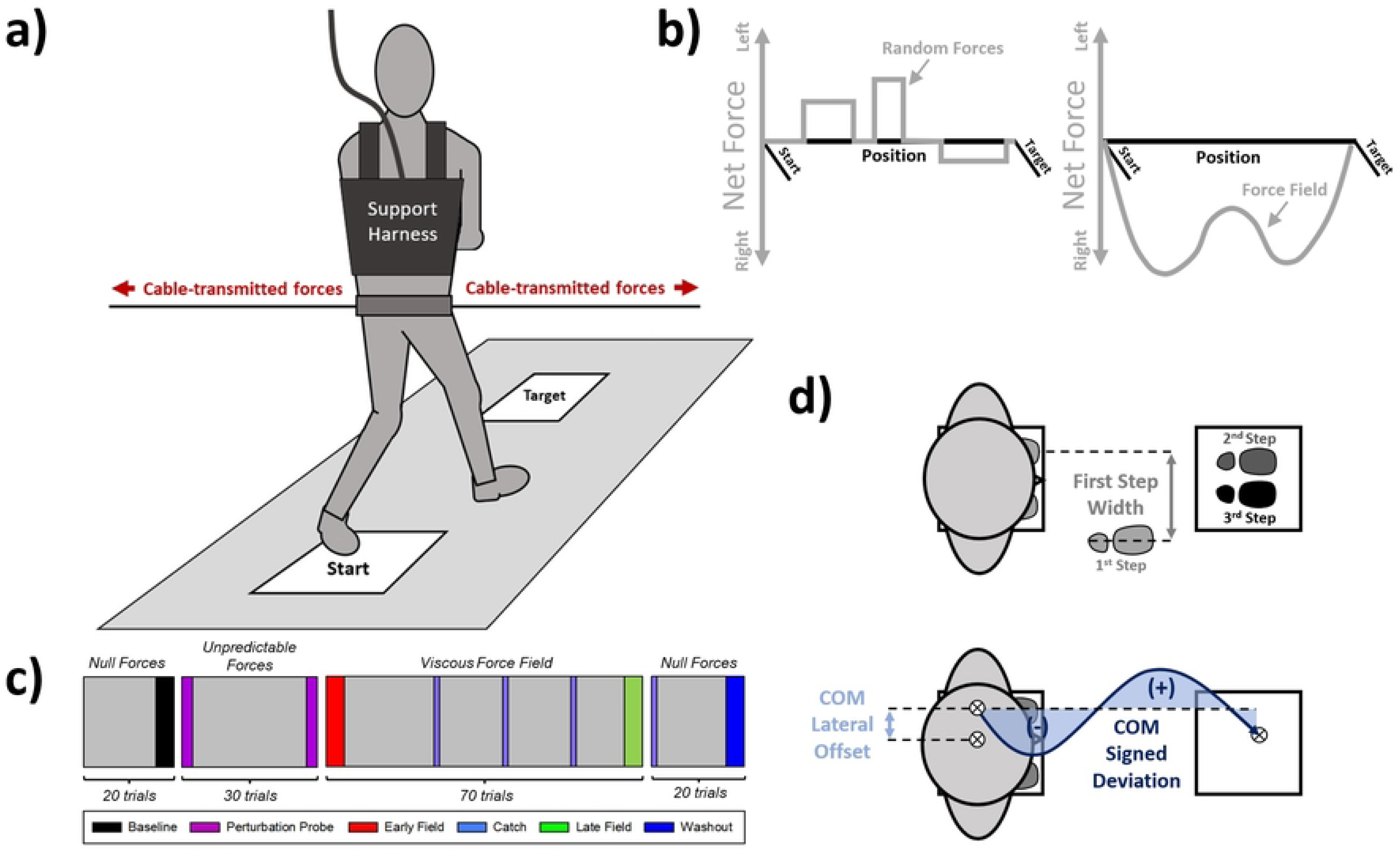
A) Experimental setup. Participants stepped forward from a 0.3 m × 0.3 m starting location to an equal-sized target location projected onto a walking surface. Participants were instructed to attempt to reach the target despite possibly experiencing lateral disruptive forces delivered by a cable-driven robotic system. Participants wore a free-moving support harness for safety in case of falls. B) Disruptive forces experienced by participants. During the Unpredictable Forces block, participants experienced a randomized series of brief force impulses (see Methods for randomization parameters). During the Force Field block, participants experienced a continuous lateral force field proportional to their forward velocity (80 N/(m/s)). Note the waveform of the Force Field reflects a standard forward velocity profile. C) Trial protocol and experiment blocking for the Perturbation Group. Colored blocks indicate trials selected for analysis. The Control Group, which were collected as part of a previous study (Bucklin et al., 2019), did not experience the Unpredictable Forces. D) Assessed metrics underlying balance control. We assessed control of COM position by calculating COM Signed Deviation, the signed area of COM trajectory. To assess underlying control strategies, we computed First Step Width and the lateral offset in the participant’s COM when initiating first toe-off.

### 2.2a Creation of External Walking Environments

During all trials, a cable-driven robotic device (the Agility Trainer; Brown, Wu, Huang, & Gordon, 2017) applied laterally directed forces to the participant’s pelvis (Fig. 1a). Participants wore a snug pelvic harness that was attached on each side to an independent cable connected to a series-elastic linear motor. The cables were routed through a trolley system that allowed free fore-aft motion of the participant while maintaining lateral forces in the cables sufficient to avoid slack in the cables and a net force of zero on the participant. Load cells (Transducer Techniques, Temecular, CA) recorded forces acting on the cable-harness attachments to provide feedback to a real-time control system in LabView (National Instruments Corporation, Austin, TX).

Depending on the trial condition, the Agility Trainer produced one of three distinct types of forces: Null Forces, Unpredictable Forces, and the Viscous Force Field.

#### Null Forces

The Agility Trainer applied zero net force to the participant (Fig. 1b).

#### Unpredictable Forces

The Agility Trainer delivered brief balance perturbations to participants by applying impulsive lateral forces that ranged in magnitude (75N to 12% body weight), direction (left or right), duration (200 to 500 ms), frequency per trial (0 to 3) and onset timing (0 to 500 ms) (Fig. 1b). These parameters were determined based on pilot testing to ensure the perturbations elicited by the forces would be perceivable and sufficient to cause lateral balance displacements without causing falls or exceeding the motor limits of the cable robot. To make the forces unpredictable to the participants, the exact values of each parameter were generated randomly for each trial.

#### Viscous Force Field

The Agility Trainer applied a continuous lateral force field to the participant’s right side (Fig. 1b) proportional in magnitude to forward walking velocity. The applied force field (80 N/(m/s)) was the same for all participants and selected to provide a walking environment that was challenging but not strong enough to evoke a fall (maximum applied force across all trials was 17.1 ± 1.1% of participant body weight). Forward walking velocity was calculated on-line using the derivative of COM position measured by a string potentiometer attached to the front of the participant’s pelvic harness. Position data was sampled at 100 Hz and low pass filtered in real time (10 Hz cut-off) to reject spikes in the derivative (velocity) due to noise. For brevity, the Viscous Force Field will hereafter be referred to as simply the *Force Field*.

Participants were never informed about what forces they would be experiencing prior to each trial. To prevent participants from attempting to “probe” which field they would receive at the beginning of each trial (e.g., by leaning slightly forward prior to first toe-off), the Force Field did not onset until participants’ forward movement exceeded a velocity threshold (0.40 m/s).

### 2.2b Forward walking velocity feedback

Visual feedback about forward velocity was provided after the completion of each trial to help participants maintain a similar forward walking velocity across all trials. A monitor positioned at the end of the walking path displayed the feedback stating either “too slow”, “too fast”, or “good job” if the participant’s peak forward velocity was respectively under, within, or above a range of 1.2 ± 0.1 m/s.

### 2.2 c Kinematic measurements

We used a 12-camera motion capture system (Qualisys, Gothenburg Sweden) recording at 100 Hz to capture 3D coordinates of reflective markers placed on the participant’s pelvis and feet. Specifically, we tracked pelvis location using three markers arranged in a triangle cluster over the S2 and T12 vertebrae and markers placed bilaterally on the greater trochanter, and we tracked the positions of each foot with markers placed on the lateral malleolus, calcaneus, and 2^nd^, 3^rd^, and 5^th^ metatarsals, resulting in a total of 15 markers used for kinematic measurements.

### 2.3 Protocol

Participants experienced slightly different trial schedules depending on whether they were in the **Control** or **Perturbation** group. Chronologically, the trials for the Control group were: 20 Baseline trials (Null Forces), then 70 Force Field trials, then 20 Washout trials (Null Forces), resulting in 110 total trials (see Fig. 1c of Bucklin et al., 2019). Three Force Field trials (45^th^, 60^th^, and 75^th^ overall trials) were replaced with Null Force trials. These trials, delivered when the participant was expecting the Force Field, served as “catch trials” to monitor any behaviors by the participant in anticipation of the Force Field.

The Perturbation group experienced an identical blocking of walking trials with one exception. Between the Baseline and Force Field trial blocks the Perturbation group experienced 30 trials with Unpredictable Forces, resulting in 140 total trials (Fig. 1c). To quantify immediate effects of the Unpredictable Forces trials on balance performance (i.e., effects prior to Force Field trials), the first and last two trials of Unpredictable Forces were prescribed to be of fixed magnitude, direction, and timing to create a controlled condition to make before vs. after comparisons. The trials were prescribed to use a single square-wave force impulse with a magnitude of 12% of the participant’s bodyweight and onset 200 ms after the start signal. The impulse was directed to the participant’s right side for the first and last trials of the Unpredictable Forces block (**Right Perturbation Probes)**, and to the participant’s left side for the second and second-to-last trials (**Left Perturbation Probes**).

### 2.4 Procedure

Each walking trial began with the participant standing motionless with both feet located in the start target. Next, participants heard an auditory “get ready” command followed by a “beep” cue. At the cue, participants stepped quickly to the end target, leading always with their right foot. The trial concluded when both feet were located within the end target. At this time, a second auditory “beep” signaled the trial was over and that the participant should return to the start target. No forces were applied to the participant as they reset their position to the start target. Participants were instructed to cross their arms during each trial to eliminate possible confounds from arm movements. Aside from these instructions, participants were told to complete the task in the manner they felt most comfortable with.

Before starting the experimental trials, participants experienced five practice trials with Null Forces to develop a basic understanding of the task and task requirements, such as becoming familiar with the desired forward velocity range. Participants then received two practice trials with the Force Field to receive a basic understanding of the general magnitude of force that would be applied to them during the Force Field trials.

### 2.5 Data processing and analysis

Kinematic marker data was processed using Visual3D (C-Motion, Germantown, MD) and a custom MATLAB (MathWorks, Natick, MA) program. Marker data was gap-filled and low-pass filtered (Butterworth, 6 Hz cut-off frequency). We identified gait events as moments of toe-off (moments when the foot leaves the ground) and heel strike (moments when the foot contacts the ground) based on the inferior-superior positions of the calcaneus and 2^nd^ metatarsal markers. Toe-off was identified per step as the local minimum of the 2^nd^ metatarsal and heel strike was identified per step as the local minimum of the calcaneal marker. All gait events were visually inspected to verify accurate event detection and were manually re-labeled as necessary. COM position was calculated in Visual3D as the center of the pelvis model, determined by the three pelvic and two greater trochanter markers.

To characterize control of COM trajectory, we analyzed kinematic data of the COM trajectory between the start and end targets, and calculated **COM signed deviation** (Fig. 1d). COM signed deviation is the signed area of the COM trajectory, assessed from the moment of first toe-off to the moment of the trial-ending “beep” cue, relative to a straight-line path originating from the COM position at first toe-off. We normalized this relative to Baseline by subtracting the averaged COM signed deviation taken over the last four Baseline trials. The COM signed deviation was thus an error metric: greater directional biases in COM trajectory (via differences between areas on either side of the straight path) led to larger COM signed deviation values and indicate greater perturbation of balance. Positive COM signed deviation values indicate leftward bias compared to baseline, and negative values indicate rightward bias compared to baseline.

To gain insight into the strategies underlying the control of COM trajectory, we also evaluated lateral movement of the COM prior to first toe-off (**COM lateral offset**) as a potential anticipatory strategy and the width of the first step with the right foot (**First Step Width**) (Fig. 1d). We calculated COM lateral offset as the lateral distance between the static COM position and the COM position at first toe-off of the right foot. First step width was calculated as the medio-lateral distance between the left calcaneus marker at the start cue and the right calcaneus marker at first right heel strike.

To assess changes in our three performance metrics (**COM signed deviation**, **COM lateral offset, and first step width**), we analyzed data from five experimental periods (Fig. 1c). The five experimental periods were: **Baseline**, **Early Field**, **Catch Trials**, **Late Field**, and **Washout**. The purpose of the Baseline period was to establish Baseline walking criteria as a reference for later performance. The purpose of the Early and Late Field periods was to assess immediate and latent effects, respectively, of the Force Field trials. The purpose of the Catch Trials, as mentioned earlier, was to assess strictly anticipatory control strategies by the participant. And the purpose of the Washout period was to determine if participants returned to Baseline walking levels (i.e., determine if there were any lasting effects of the Force Field). Values for each period were calculated as an average of values from four specific trials within that period. For Baseline, we used the last four Baseline trials; for Early Field, we used the first four Force Field trials; for Catch Trials, we used the three catch trials and the first Washout trial (which served as an additional *de facto* Catch Trial); for Late Field, we used the last four Force Field trials; and for Washout, we used the last four Washout trials.

For the Perturbation group, we assessed COM signed deviation in the **Left and Right Perturbation Probes** to assess immediate effects (before vs. after) of the Unpredictable Forces block.

### 2.6 Statistical Analysis

To evaluate participant adaptation to the Force Field, we used mixed ANOVAs with a repeated measures factor of experimental period (*Baseline, Early Field, Late Field,* and *Washout*) and a between-subject factor of group (*Control* and *Perturbation*) to evaluate COM signed deviation, COM lateral offset, and first step width. We then evaluated the effects of Catch trials using a pair of mixed ANOVAs. The first ANOVA employed a repeated measures factor of experimental period (*Baseline* and *Catch Trial*) to evaluate how participants developed anticipatory strategies in response to the Force Field that would bias their walking in comparison to Baseline trials. The second ANOVA employed a repeated measures factor comparing *Catch* trials to the trials immediately preceding each catch trial (*Pre-Catch*) to evaluate how anticipatory strategies would bias their walking in comparison to Force Field trials.

Greenhouse-Geisser corrections were used where the assumption of sphericity was violated. When the assumption of equal variances was violated for between-subjects comparisons, unpaired comparisons with equal variances not assumed were performed instead. Holm-corrected post-hoc tests and simple effects analyses were run to follow up on significant effects.

## 3. Results

All participants complied with the experimental instructions and were able to perform the experiment with general success and without falls, reported fatigue, or reported discomfort. One participant used a large stride (leaping from the start to target locations) to complete most of the trials, which was different from the strategy used by all other participants. This led to her data deviating greatly from the other participants’ data. Therefore, we excluded her data from the subsequent statistics and analysis.

Our statistical tests revealed many main effects and interaction effects for our analyses which, due to their extent, would entail a considerable amount of space to comprehensively report. Therefore, for the sake of conciseness, we present only the highest order effects for each analysis, while providing complete findings in the supplementary material.

### 3.1 Unpredictable Forces

As a design check, we first determined if the forces delivered in the Unpredictable Forces block successfully disrupted the balance of participants in the Perturbation group. Observation of participants during the experiment as well as visual inspection of COM signed deviation data indicated that participants exhibited large bilateral deviations of their COM trajectory. To confirm this observation, we compared the mean of the absolute (i.e., unsigned) COM deviations and the variability (SD) of COM signed deviations across Unpredictable Forces trials versus across Baseline trials. We found that overall absolute COM deviations were greater in the Unpredictable Forces block (0.037 ± 0.010 m^2^) than Baseline (0.013 ± 0.005 m^2^), *t*(11) = 7.10, *p* < .001, *d* = 2.05. The SD of COM signed deviation was likewise greater in the Unpredictable Forces block (0.045 ± 0.014 m^2^) than the Baseline block (0.016 ± 0.006 m^2^), *t*(11) = 6.57, *d* = 1.90 (see Fig. 2). Thus, we concluded that participants’ walking trajectory was successfully perturbed by the Unpredictable Forces.

**Figure 2.**
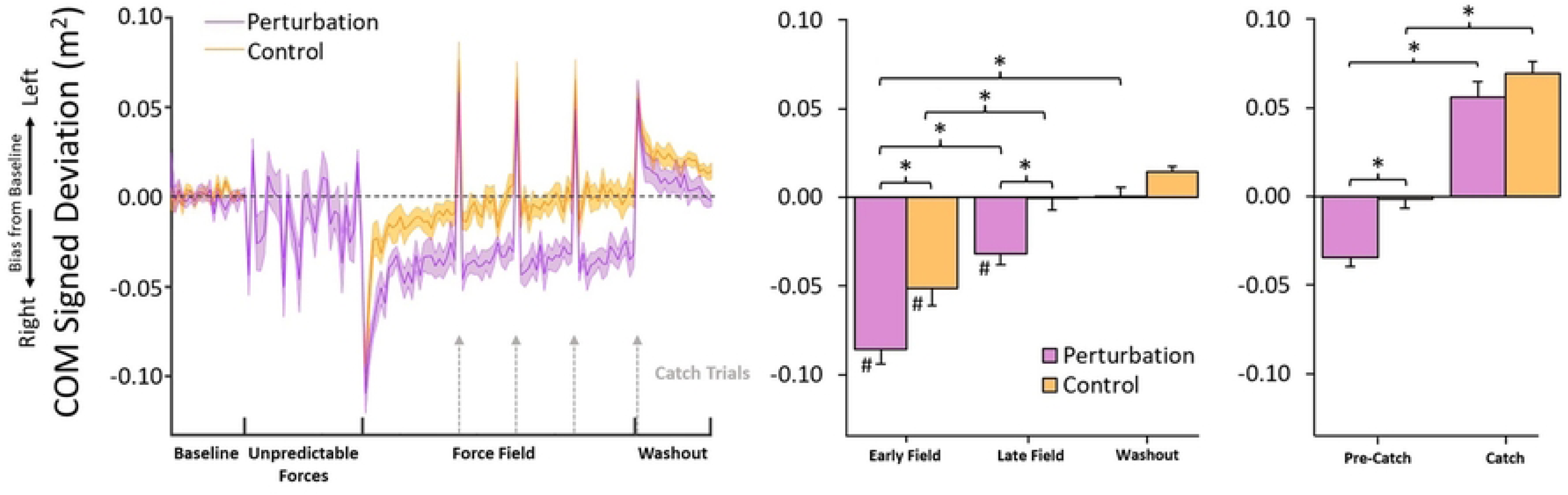
COM signed deviation results. During the Force Field trials, the Perturbation Group showed larger initial errors (rightward bias in COM signed deviation due to the rightward Force Field), slower decreases in error, and larger remaining errors following the Force Field Trials when compared to the Control Group. Both groups showed similar large errors on Catch trials.

We next investigated if participants adapted to the Unpredictable Forces over time. Specifically, we investigated if participants improved performance (i.e., reduced the magnitude of their COM signed deviation) by the end of the Unpredictable Forces block. Assessment of COM signed deviation during the Left and Right Perturbation Probes found that participants slightly improved performance from the first to last Right Perturbation Probes (−0.04 ± 0.04 m^2^ and −0.02 ± 0.02 m^2^, respectively), *t*(11) = 1.86, *p* < .05, *d* = 0.54, but not from the first to last Left Perturbation Probes, (0.03 ± 0.02 m^2^ and 0.02 ± 0.02 m^2^, respectively), *p* > .2. We interpreted this as participants showing a slight adaptation to the Unpredictable Forces, but not a substantial adaptation, as their strategy was only moderately effective at reducing balance errors for Right Perturbation Probes and not effective at reducing balance errors for Left Perturbation Probes.

### 3.2 Adaptation to Force Field

The central questions of the study concerned how participants adapted to the Force Field and if these adaptations differed between the Control group and Perturbation group. As stated earlier, we considered two competing hypotheses: that the Unpredictable Forces would inhibit predictive control by producing lasting reductions in error sensitivity (**Hypothesis 1**), or that Unpredictable Forces would facilitate predictive control by prompting protective strategies that stabilize initial movements and reduce initial errors (**Hypothesis 2**).

### 3.2 a Center of Mass Trajectories

Fig. 2’s left panel shows average COM signed deviation (normalized to baseline values) for the Perturbation and Control groups during the experiment, and the right panel shows the averages at each experimental period. During initial exposure to the Force Field (Early Field), both groups displayed significant lateral biases in COM deviation towards the right (negative COM signed deviations), aligned with the Force Field’s direction. Practice with the Force Field reduced the magnitude of the deviations, approaching baseline levels by the end of the training. When the Force Field was removed (Catch and Washout trials), both groups exhibited large lateral COM deviations towards the left (positive COM signed deviations) as a result of overcompensating for the expected but absent Force Field (see section **3.3 Catch Trials**). These deviations decreased as participants re-adjusted to Null Forces during washout trials. Overall, the Perturbation group appeared to exhibit larger deviations than the Control group at each stage of the Force Field trials.

Mixed ANOVA indicated a significant interaction between Period and Group, *F*(1.90,43.74) = 3.71, *p* < .05, *η*_p_ ^2^ = 0.14. Simple effects analysis indicated that COM signed deviation changed as a function of Period for both the Perturbation group, *F*(1.77,19.42) = 47.63, *p* < .001, *η*_p_ ^2^ = 0.81, and the Control group, *F*(1.84,22.12) = 24.44, *p* < .001, *η*_p_ ^2^ = 0.67. For the Perturbation group, COM signed deviation showed the most rightward bias in the Early Field (−0.08 ± 0.03 m^2^), which was different from baseline, *p* < .001, *d* = 4.20. Early Field COM signed deviation was also greater than in the Late Field (−0.03 ± 0.02 m^2^), *p* < .001, *d* = 2.65. Late Field COM signed deviation was greater than baseline, *p* < .01, *d* = 1.55, indicating that participants were not able to adapt balance performance to baseline levels before the end of the Force Field trials. There was no difference between baseline COM signed deviation and washout (0.00 ± 0.02 m^2^), *p* > .9.

For the Control group, COM signed deviation also showed the most rightward bias in the Early Field (−0.05 ± 0.03 m^2^), which was different from baseline, *p* < .001, *d* = 2.32. Early Field COM signed deviation in the Early Field was also, like the Perturbation group, greater than in the Late Field (0.00 ± 0.03 m^2^), *p* < .001, *d* = 2.31. Unlike the Perturbation group, however, Late Field COM signed deviation was not different from baseline, *p* > .9, indicating that the Control group returned balance performance to baseline levels before the end of the Force Field trials. There was no difference in COM signed deviation between Late Field and washout (0.01 ± 0.01 m^2^), nor baseline and washout, both *p* > .2.

Simple effects analysis finally indicated that the main effect of Group persisted at each individual experimental Period except baseline (where both groups were normalized to zero); the Perturbation group in the Early Field exhibited greater rightward bias in COM signed deviation than the Control group, *p* < .05, *d* = 1.06, and also exhibited greater rightward bias in the Late Field than the Control group, *p* < .01, *d* = 1.31. Interestingly, the Perturbation group also exhibited greater rightward bias in COM signed deviation in the washout period than the Control group, *p <* .05, *d* = 0.99.

To summarize, the COM signed deviation results indicated that both groups experienced substantial balance disruptions at the start of the Force Field trials but adapted to the Force Field and improved their walking performance with practice. However, the Perturbation Group exhibited worse initial performance in the Force Field, slower adaptation to the Force Field, and (*ipso facto*) worse overall performance with the Force Field than the Control Group. This supports **Hypothesis 1** – that prior exposure to the Unpredictable Forces inhibited the adaptation of predictive control strategies to counter the consistent conditions of the Force Field trials.

### 3.2b Center of Mass Lateral Offset

Fig. 3’s left panel shows average COM lateral offset for the Perturbation and Control groups, and the right panel shows the averages at each experimental period. Lateral offset was positive across all trials for all participants, indicating that COM always shifted toward the left limb prior to the first step’s toe-off (which was always performed with the right limb). Based on observations during experiments and visual inspection of lateral offset data, when first exposed to the Force Field (Early Field), both groups exhibited relatively little change, but perhaps a slight increase, in lateral offset. With experience walking in the Force Field, lateral offset increased substantially for the Control group but not for the Perturbation group. When the Force Field was removed (i.e., during washout trials), lateral offset for both groups appeared to trend back towards values similar to baseline.

**Figure 3.**
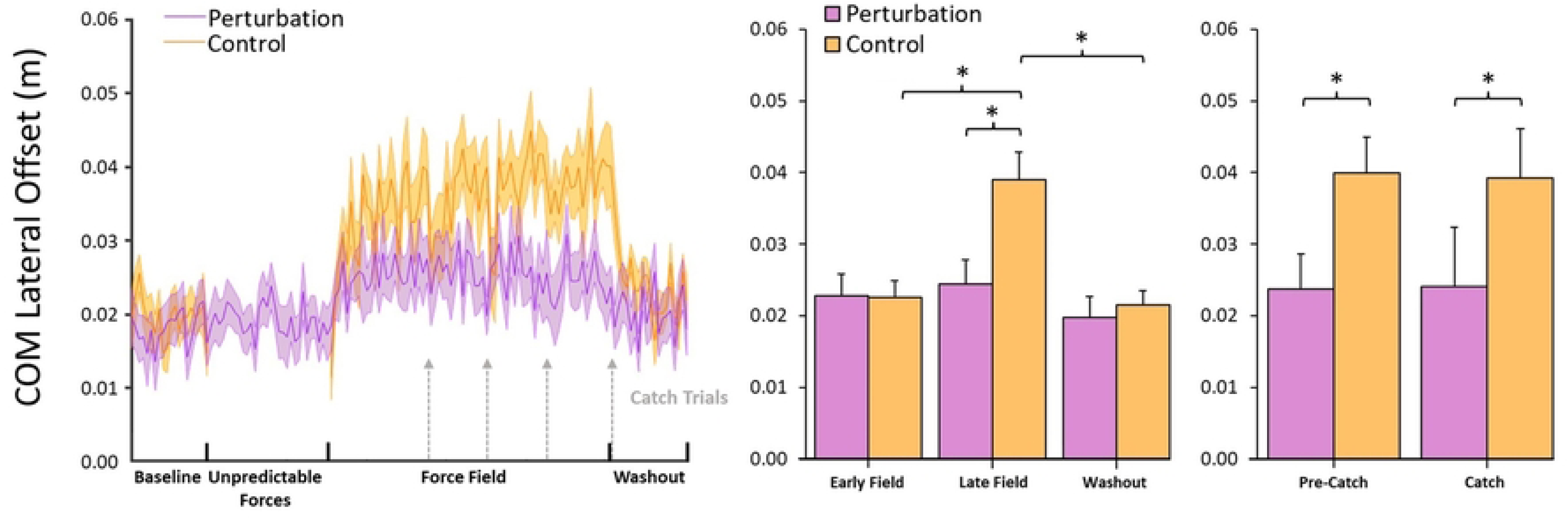
COM lateral offset results. Participants in the Perturbation group did not exhibit changes in COM lateral offset during the Unpredictable Forces trials. During the Force Field trials, Control participants gradually modulated lateral offset further to the left, but participants in the Perturbation group never significantly modulated their lateral offset differently from baseline. Lateral offset was unaffected by Catch trials in both groups.

A mixed ANOVA indicated a significant interaction between experimental Period and group on lateral offset, *F*(2.02,46.38) = 6.61, *p* < .01, *η*_p_ ^2^ = 0.22. Simple effects analysis revealed that lateral offset did not change as a function of Period for the Perturbation group, *p* > .05, but did change as a function of Period for the Control group, *F*(1.81,21.69) = 15.58, *p* < .001, *η*_p_ ^2^ = 0.57. For the Control group, lateral offset was not different between the baseline (0.02 ± 0.01 m), Early Field (0.02 ± 0.01 m), or washout (0.02 ± 0.01 m) trials, all *p* > .7. Lateral offset in the Late Field trials (0.04 ± 0.01 m), however, was greater than every other period (baseline, *p* < .001, *d* = 2.09; Early Field, *p* < .001, *d* = 1.71; washout, *p* < .001, *d* = 1.82). Likewise, lateral offset was not different between groups in the baseline (Control group = 0.02 ± 0.01 m), Early Field (Control group = 0.02 ± 0.01 m), or washout (Control group = 0.02 ± 0.01 m) trials, all *p* > .6, but was greater for the Control group than the Perturbation group (Control group = 0.02 ± 0.01 m) in the Late Field trials, *p* < .01, *d* = 1.13.

To summarize, these results indicate that while Control participants did not initially modulate lateral offset during Force Field trials, they eventually adopted a strategy of offsetting their COM further to the right after continued practice with Force Field trials. Participants in the Perturbation group, however, never significantly modulated their lateral offset differently from baseline. As COM lateral offset is specifically a predictive control strategy for countering expected lateral balance disruptions, these results support **Hypothesis 1**.

### 3.2c First Step Width

Fig. 4’s top panel shows group averages for starting left foot position and first right step position (the two reference points for calculating first step width), and the bottom panel shows the actual first step width values normalized to baseline. Visual inspection of the data and observations of participants during the experiment suggested that as participants first experienced the Force Field, they began to adopt wider first steps. However, those in the Control Group immediately (i.e., within the first few trials) began taking much wider first steps than those in the Perturbation Group, and the width of these steps increased only marginally over the course of the Force Field trials. The Perturbation Group, in contrast, only marginally increased their first step width at first, but substantially increased this width over the course of the Force Field trials.

**Figure 4.**
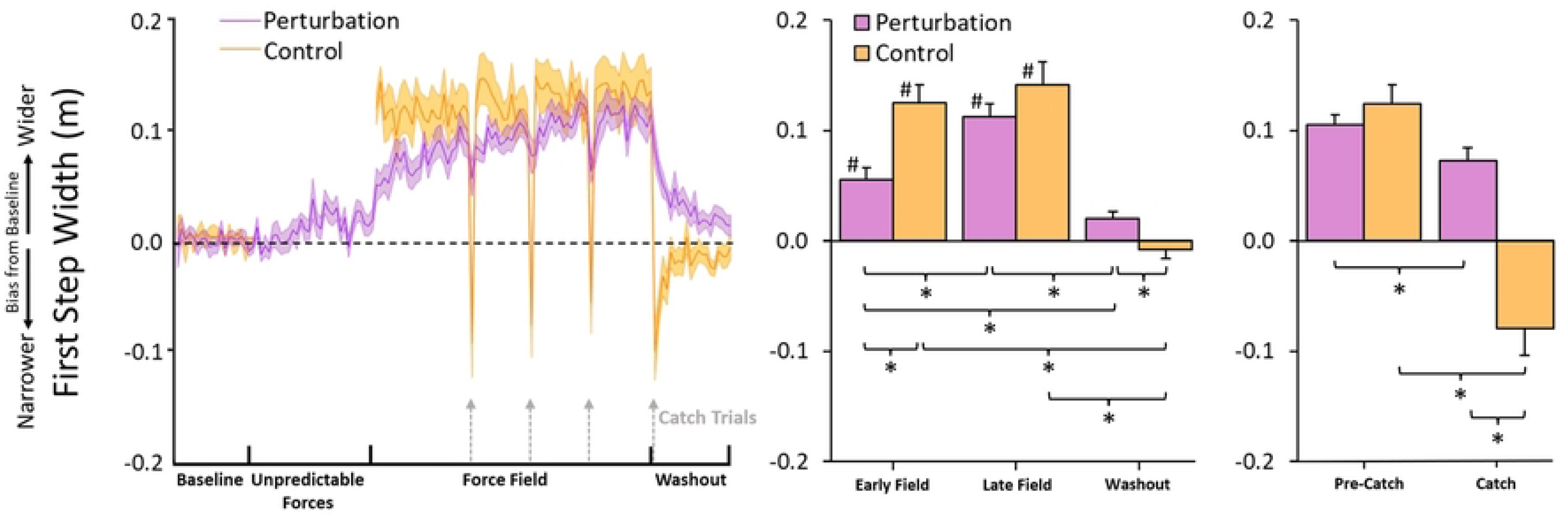
First step width results. Both groups adopted a strategy of widening their first step to counter the effects of the Force Field, but the Control group’s response was more immediate and more pronounced, while the Perturbation group’s response was more gradual and less pronounced. Participants in both groups exhibited decreases in first step width during Catch trials as a reaction to the unexpected absence of the Force Field, but these decreases were substantially larger for the Control group.

A mixed ANOVA indicated a significant interaction between Period and Group, *F*(1.74,40.00) = 9.10, *p* < .001, *η*_p_ ^2^ = .78. Simple effects analysis confirmed that first step width changed as a function of Period for both the Perturbation group, *F*(3,33) = 46.24, *p* < .001, *η*_p_ ^2^ = 0.81, and the Control group, *F*(1.61,19.26) = 47.54, *p* < .001, *η*_p_ ^2^ = 0.80. For the Perturbation group, first step width increased from baseline to Early Field (0.06 ± 0.04 m), *p* < .001, *d* = 1.82, and from Early Field to Late Field (0.11 ± 0.04 m), *p* < .001, *d* = 1.90. First step width in the Late Field was also greater than baseline, *p* < .001, *d* = 3.72. First step width then decreased from Late Field to washout (0.02 ± 0.02 m), *p* < .001, *d* = 3.06. First step width in the washout trials was also lower than in the Early Field, *p* < .01, *d* = 1.16. There was no difference between first step width in washout and baseline trials, *p* > .05, indicating that participants in the Perturbation group returned first step width back to baseline-similar levels by the end of the washout trials. For the Control group, first step width likewise increased from baseline to Early Field (0.13 ± 0.06 m), *p* < .001, *d* = 2.52. However, unlike the Perturbation group, first step width in the Control group did not increase from Early Field to Late Field, *p* > .6. First step width in the Late Field was, like the Early Field, greater than baseline, *p* < .001, *d* = 2.85, and decreased from the Late Field to washout (−0.01 ± 0.03), *p* < .001, *d* = 3.01. There was no difference between first step width in the baseline and washout trials, *p* > .6, indicating that participants in the Control group also returned first step width back to baseline-similar levels by the end of the washout trials.

Simple effects analysis finally indicated that between-group differences in first step width varied with experimental Period. The Control group exhibited larger first step width than the Perturbation group during Early Field trials, *p* < .01, *d* = 1.40, but exhibited smaller first step width than the Perturbation group in washout trials, *p* < .05, *d* = 1.08. There were no differences between groups at baseline (as both groups were normalized to zero), *p* > .4, or Late Field, *p* > .2.

To summarize, these results indicated that while both groups adopted a strategy of widening their first step to counter the effects of the Force Field, the Control group’s adaptation was more immediate and more pronounced, while the Perturbation group’s response was more gradual and less pronounced. These results further support **Hypothesis 1**.

### 3.3 Catch Trials

During catch trials and early washout trials, the unexpected absence of the Force Field resulted in the predictive movements that participants adapted to counter the Force Field producing large lateral COM deviations toward their left (positive COM signed deviation values) as well as large changes in their step placements.

### 3.3a Center of Mass Trajectories

A mixed ANOVA confirmed a main effect of Pre-Catch vs. Catch Trials, *F*(1,23) = 192.72, *p* < .001, *η*_p_ ^2^ = 0.89. Lateral COM deviations changed significantly from Pre-Catch trials (−0.02 ± 0.00 *m^2^*) to Catch trials (0.06 ± 0.03 m^2^). There was also a main effect of Group, *F*(1,23) = 9.62, *p* <.01, *η*_p_ ^2^ = 0.30, indicating that the Control group (0.03 ± 0.04 m^2^) exhibited more leftward bias in COM signed deviation over both trials than the Perturbation group (0.01 ± 0.05 m^2^). A Group × Trial interaction was not significant, *p* > .1. While the Group differences suggested a differential effect of the Force Field on COM Trajectories, the lack of an interaction (i.e., lack of difference between how the catch trials affected each group) means that neither **Hypothesis 1** nor **Hypothesis 2** are supported or rejected by these results.

### 3.3b COM Lateral Offset

A mixed ANOVA indicated no effect of Trial on COM lateral offset, *p* > .9, and likewise no interaction between Group and Trial, *p* > .7. This was to be expected, as COM lateral offset occurs entirely before the onset of the Force Field and thus would not be affect by the presence or absence of it. The ANOVA confirmed only the main effect of group, *F*(1,23) = 10.86, *p* < .01, *η*_p_ ^2^ = 0.32, that was previously discussed in section **3.2b**; The Control group (0.04 ± 0.01 m) adopted a larger bias in their lateral offset than the Perturbation group (0.02 ± 0.0 m), indicating greater adaptation of predictive control strategies. Thus, like COM trajectories during catch trials, these results cannot confirm or reject either **Hypothesis 1** or **Hypothesis 2**.

### 3.3c First Step Width

A mixed ANOVA indicated an interaction between Trial and Group, *F*(1,23) = 70.92, *p* < .001, *η* ^2^ = 0.76. Simple effects analysis revealed that there was no difference in first step width between Group for Pre-Catch trials, *p* > .3. There was, however, a difference between Group during Catch trials, *p* < .001, *d* = 2.15; the Control group (−0.08 ± 0.09 m) exhibited much narrower first step widths on Catch trials than the Perturbation group (0.07 ± 0.05 m), indicating that their first step was much more affected by the catch trial manipulation. This was confirmed by the simple effects of Trial, which indicated that while first step width decreased as a function of Trial for both the Control group, *p* < .001, *d* = 3.29, and the Perturbation group, *p* < .001, *d* = 0.95, the greater effect size for the Control group indicated that first step width decreased more during the Catch trials.

In sum, these results support **Hypothesis 1** by indicating that participants in the Control group adopted a much stronger predictive control strategy (modulating first step width) than those in the Perturbation group, which led to greater errors during Catch trials.

## 4. Discussion

The current study investigated if exposing participants to environmental uncertainty via unpredictable balance perturbations impeded their adaptation of predictive control strategies to a novel but consistent viscous force field. To investigate this, the current study assessed two competing hypotheses. **Hypothesis 1** was that prior exposure to environmental uncertainty would impede the adaptation of predictive control strategies by prompting participants to instead adapt protective strategies that reduce attention to motor errors and thereby slow trial-to-trial movement corrections. **Hypothesis 2** was that prior exposure to environmental uncertainty would lead to the adaptation of control strategies that would stabilize movements during early exposure to the force field and thereby enhance adaptation to the force field. Our results overall consistently supported **Hypothesis 1** over **Hypothesis 2**. Participants in the Perturbation group exhibited less prominent predictive control strategies throughout the force field trials, were less affected by the catch trials, and reduced their kinematic errors (i.e., deviations in their COM trajectories) less than the Control group, suggesting that prior exposure to Unpredictable Forces trials inhibited their development of predictive control strategies.

### 4.1 Prior Environmental Uncertainty Inhibited Adaptation of Error Reduction Strategies to Offset a Consistent Environment

Both groups of participants experienced large disruptions to their balance when first experiencing the Force Field as evidenced by large rightward biases in their COM trajectories, i.e., large (negative) COM signed deviations. With practice, both groups adapted to the Force Field and the deviations in the COM trajectories decreased. However, the Control group exhibited much more immediate reductions in COM signed deviation and greater reductions by the end of training than the Perturbation group. In fact, the Control group was able to reduce COM signed deviations to baseline-similar values by the end of the Force Field trials while the Perturbation group was not. On Catch trials, both groups of participants were caught off guard by the unexpected removal of the Force Field and exhibited large deviations in the COM trajectory in the opposite direction of the anticipated Force Field (i.e., to the left). This indicates that both groups were employing, to some degree, a predictive strategy to resist the rightward influence of the Force Field (see discussion about COM Lateral Offset below for elaboration) that led to overcorrections in balance control when the Force Field was removed. This change due to Catch trials was not different between groups, but the greater overall adaptation in COM signed deviation in the Control group nevertheless indicate that the Control group was less inhibited during adaptation than the Perturbation group.

Participants in the Control group likewise exhibited larger COM lateral offset than the Perturbation group in the Late Field trials, which indicates the stronger adaptation of predictive control. Finally, participants in the Control group adopted larger, more immediate adjustments in first step width than those in the Perturbation group and these adjustments were more affected by Catch trials in the Control group than they were for the Perturbation group.

These results offer compelling evidence that overall adaptation to a consistent external walking environment is inhibited by prior exposure to an uncertain external walking environment. This follows similar findings of sensorimotor uncertainty inhibiting error-based learning across a variety of task domains, such as reaching (Albert et al., 2021; Herzfeld et al., 2014; Therrien, Wolpert, & Bastian; 2018), visuomotor rotations (Fernandes, Stevenson, & Körding, 2012) and saccadic fixations (Havermann & Lappe, 2010). In these studies, variability is introduced to perturbations or sensorimotor disruptions imposed on the participant. When this variability is increased to levels that are intractable to the participants (i.e., cannot be predicted), their adaptation rates decrease and they display persistent, residual motor errors, even when the variability is removed again. That this phenomenon persists across varying task domains indicates a general principle that exposure to task uncertainty produces lingering effects on subsequent motor adaptations.

### 4.2 Prior Environmental Uncertainty Inhibited Predictive Control Strategies

Bucklin et al. (2019) found that participants adopted a strategy of shifting COM lateral offset further than normal as a way of countering the Force Field by pre-emptively biasing their COM trajectory in the opposite direction (i.e., they initiated gait with their right leg and therefore shifted their COM further to the left in anticipation of the Force Field’s rightward forces). The adaptation of this predictive strategy was gradual, without much change in the early Force Field trials but showing a clearly emergent bias by the end of the Force Field trials.

The current study found that, after exposing participants first to the Unpredictable Force trials, those participants did not adopt any changes to their COM lateral offset over the course of the Force Field trials.

Bucklin et al. (2019) also observed that participants adopted a wider first step with their right foot as a way of creating a wider base of support to resist against the rightward Force Field. They adopted this strategy immediately, showing substantially wider first steps within the first few Force Field trials, and then marginally widened their steps even further over the remaining trials. During Catch trials, the unexpected removal of the Force Field required participants to reactively plant a much narrower first step to attempt to catch themselves as they fell leftward (due to their COM lateral offset biasing their COM trajectory so strongly to the left). For Control participants, this first step width during Catch trials was narrower than baseline and was frequently even negative as their first step was placed further to the left than their left limb in stance phase (i.e., cross-stepping).

The current study found that, after exposing participants first to the Unpredictable Force trials, those participants did not immediately adopt wider first steps during Force Field trials like the Control group, but rather they gradually increased their first step width over the course of Force Field trials. Moreover, the first step width of Perturbation group participants was not nearly as affected by Catch trials. During Catch trials, participants exhibited first steps that were narrower than the prior step but remained wider than they were at Baseline, unlike the Control group.

These results offer evidence that the inhibition of overall adaptation to a consistent external walking environment by prior exposure to an uncertain external walking environment, as highlighted in the previous section, is driven by the inhibition of *predictive* control strategies in particular.

Based on these findings, it is possible that discrete stepping maneuvers follow a Bayesian learning framework. For the control of brief, discrete actions (such as the Force Field stepping task used in the current study), when encountering task situations with high amounts of uncertainty, humans tend to shift reliance away from sensory feedback and more towards fixed expectations about the task (Chambers et al., 2018; Darlington et al., 2018; Jarbo et al., 2018; Palmer et al., 2019). This study and others (Whittier, Weller, & Fling, 2022) suggest this behavior extends to balance-related discrete stepping tasks. Discrete stepping could follow a process of Bayesian inference for learning and adaptation, where feedforward control of movements underlying the stepping process (COM lateral offset, first step placement, etc.) is tuned both by sensorimotor feedback and learned expectations about the movement environment (Körding & Wolpert, 2004). Such a framework would clarify the role of environmental uncertainty during discrete locomotor adaptation and could be pursued with a more precise account of the relative weighting of sensorimotor feedback versus learned expectations on underlying stepping movements, such as with the methodology outlined by Whittier, Weller, and Fling (2022).

### 4.3 Why Participants Showed Little Adaptation to the Environmental Uncertainty

A central expectation of the study was that participants in the Perturbation group would adapt to the Unpredictable Forces trials in some way that would either inhibit or enhance their subsequent ability to adapt to the Force Field trials. We were surprised, then, to observe that participants exhibited barely any reduction in kinematic errors (i.e., little reduction in COM signed deviation) from the start to the end of the Unpredictable Forces trials. One possible explanation for this is that the period of Unpredictable Forces (30 trials) was simply too short, and with longer exposure a pattern of adaptation would start to emerge. This possibility is supported by our prior work that found people exhibited reductions in kinematic errors when walking in Force Fields that unpredictably changed direction from trial to trial over 70 practice trials (Bucklin et al., 2023). However, the unpredictable characteristics (direction only) of the applied forces in this previous study were less complex than the Unpredictable Forces used in the current study. Thus, a second possibility is that it was not feasible for participants to adapt a movement strategy that could consistently reduce kinematic errors across all randomized possibilities of direction, timing, and frequency of forces. If this was the case, the Unpredictable Forces trials could have impacted participants purely through their expectations of the walking environment—by conditioning them to expect an unpredictable environment, they became hesitant to pursue the adaptation of predictive strategies.

### 4.4 Limitations

During Force Field trials, the force field would usually onset some time prior to the heel strike of the participant’s first step. This means that, in theory, the modulation of first step width as a movement strategy could have carried a reactive component alongside a predictive component. The current study was not able to investigate the degree to which participants’ first step placement was predictive or reactive, but future studies may be able to investigate this by ensuring that force field onset occurs too late for participants to be able to apply any reactive adjustments to their first step placement.

We were unable to make direct comparisons between walking behavior between the Unpredictable Forces trials and the Force Field trials because the participants experienced qualitatively different forces in the Unpredictable Forces block (random square-wave force impulses) versus the Force Field block (a viscous force field) as well as randomly different forces between trials within the Unpredictable Forces block. We have considered other ways of manipulating the predictability of walking environment conditions while preserving the essential quality of forces being experienced, such as manipulating the frequency at which the forces occur or the rate at which the force field switches direction. Doing so would allow for direct comparison between predictable vs. unpredictable trial blocks, and we are pursuing this investigation currently.

## Notes

### Competing Interest Statement

The authors have declared no competing interest.

